# Design, optimization, and inference of biphasic decay of infectious virus particles

**DOI:** 10.1101/2024.02.23.581735

**Authors:** Jérémy Seurat, Krista R. Gerbino, Justin R. Meyer, Joshua M. Borin, Joshua S. Weitz

## Abstract

Virus population dynamics are driven by counter-balancing forces of production and loss. Whereas viral production arises from complex interactions with susceptible hosts, the loss of infectious virus particles is often approximated as a first-order kinetic process. As such, experimental protocols to measure infectious virus loss are not typically designed to identify non-exponential decay processes. Here, we propose methods to evaluate if an experimental design is adequate to identify multiphasic virus particle decay and to optimize the sampling times of decay experiments, accounting for uncertainties in viral kinetics. First, we evaluate synthetic scenarios of biphasic decays, with varying decay rates and initial proportions of subpopulations. We show that robust inference of multiphasic decay is more likely when the faster decaying subpopulation predominates insofar as early samples are taken to resolve the faster decay rate. Moreover, design optimization involving non-equal spacing between observations increases the precision of estimation while reducing the number of samples. We then apply these methods to infer multiple decay rates associated with the decay of bacteriophage (‘phage’) ΦD9, an evolved isolate derived from phage Φ21. A pilot experiment confirmed that ΦD9 decay is multiphasic, but was unable to resolve the rate or proportion of the fast decaying subpopulation(s). We then applied a Fisher information matrix-based design optimization method to propose non-equally spaced sampling times. Using this strategy, we were able to robustly estimate multiple decay rates and the size of the respective subpopulations. Notably, we conclude that the vast majority (94%) of the phage ΦD9 population decays at a rate 16-fold higher than the slow decaying population. Altogether, these results provide both a rationale and a practical approach to quantitatively estimate heterogeneity in viral decay.

## 1. INTRODUCTION

The life cycle of viruses includes a balance between production and decay. Although production of virus particles from infected hosts is often the priority of environmental and health-associated studies, the numerical abundance of viruses in an environment is also shaped by the rate at which virus particles (i.e., virions) lose their infectivity over time. Mechanistically, virions can lose infectivity due to multiple processes including thermal-induced destabilization, aggregation, physical decay (e.g., the separation of tail fibers from the head), and/or damage to the viral genome (e.g., via exposure to UV light). The aggregate impact of these processes is conventionally modeled as a single, kinetic rate [7]. The decay rate can be considered as the rate constant associated with the exponential decay of infectious virions over time and is often strongly linked to temperature [7, 4]. However, virus populations need not decay exponentially [5, 8, 22].

A virus decay curve is a monotonically decreasing function over time, and the simplest exception to that of pure exponential decay is termed ‘multiphasic decay’. A population exhibiting multiphasic decay would be characterized by a set of different exponential rate constants. Hence, each subpopulation exhibits pure exponential decay but the total population does not. Multiphasic decay could arise because the population is comprised of different viral ‘types’ (e.g., species or alternatively self-assembled particle morphs) or because the process of decay involves different routes to failure. Irrespective of the underlying mechanism, inferring multiphasic decay presents unique challenges to experimental design. In general, in the case of biphasic decay, a biexponential model is used to describe the data by estimating the two slopes corresponding to each decay rate. As such, conventional approaches to equitemporal spacing of measurements may fail to resolve the differences between rate constants and the magnitude of the subpopulation corresponding to each of the different decay rates.

The design choice for viral decay experiments is therefore crucial. In this context, the study design includes the number of measurements and designation of the time of each measurement. Frequently, the design is chosen in an empirical way, for instance with equi-spaced sampling times. A wrong design choice results in poor parameter precision. For instance, the first and fastest slope may be incorrectly estimated if early time samples are not collected. Also, in the case of short studies, the slowest slope, corresponding to the more stable subpopulation, cannot be observed if there has not been enough measurements of decay.

Approaching the inference problem for multiphasic decay requires evaluating and optimizing the design elements, i.e., the duration time of the study as well as the number and the allocation of the measurement times. The problem of design choice can be solved by choosing the experimental design appropriately. Optimal design [1, 19] has shown good performance in decreasing the bias and uncertainty in parameter estimation in multiple contexts, including in virus studies [14, 21]. However, experimentalists often have limited prior information on parameters associated with viral decay rates and the relative proportion of each decay type in a population. In this case, robust design optimization accounting for these uncertainties may be required [13]. The design objective is to select the number of replicates, the number of samples, the sampling times, and (potentially) the initial viral density to accurately measure viral decay rates and associated subpopulation sizes.

Here, we use a Fisher information matrix-based design optimization method to precisely infer rates and magnitudes of subpopulations for multiphasic viral decay experiments. This enables us to investigate the importance of the viral parameters (decay rates, initial proportions of each subpopulation) when choosing the design and to optimize experimental design protocols, even in the case of parameter uncertainties. By leveraging *in silico* studies, we demonstrate a proof of principle for robust design and then apply these methods to phage ΦD9, showing that we are able to successfully infer multiphasic decay in practice. This inference approach identified a hidden majority of fast decaying phage and a subpopulation of slower decaying phage, raising new questions on the biophysical mechanisms and eco-evolutionary dynamics associated with variation in phage loss rates.

## 2. MATERIALS AND METHODS

### 2.1 Optimal design for time-series experiments

#### 2.1.1 Statistical model

A nonlinear model to describe viral density observations *y* (of length *n*) can be written as follows (eq. 1):

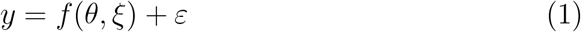

The function *f* (the structural model), depends on *θ*, the vector of the parameters describing the viral kinetics, and *ξ*, the design, provides a *n*vector of model predicted values. The *n*-vector of errors is denoted *ε*, with *ε* ∼ *N* (0, (*σ × f* (*θ, ξ*))^2^), i.e., assuming a proportional error to the structural model *f*, with *σ* the standard deviation of proportional error terms. The vector of parameters, of size *P*, to be estimated is composed of *θ* and *σ*.

#### 2.1.2 Fisher information matrix based design evaluation and optimization

A design *ξ* is defined by the number of samples *n* and their allocation in time (*t*_1_,…,*t*_*n*_), and some additional design variables (e.g., the initial viral density *V*_0_ which can otherwise be defined as a parameter).

Optimizing the design in our case is optimizing the choice of samples (number, time allocation) to maximize precision of parameter estimation. According to the Rao-Cramer inequality, the expected Fisher information matrix (FIM or *M*_*F*_ (*θ, ξ*)) is the inverse of the lower bound of the variance-covariance matrix of any unbiased estimated parameters. More details about FIM evaluation for nonlinear models are given in [1, 12, 24]. Therefore, the square root of the inverse of diagonal elements of the FIM is used to provide expected standard errors (SE) of parameters.

The D-optimality criterion Φ_*D*_ (eq. 2), widely used in the field of optimal design, consists in the determinant of the FIM normalized by *P* (the number of model parameters to be estimated) [1]:

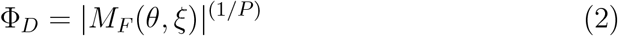

Optimization algorithms aim to find the design (given the constraints) which maximizes this criterion, i.e., which gives the best overall expected precision of estimated parameters.

The D-optimality criterion is used to perform local design evaluation or optimization, i.e., considering the model and parameter as known. Alternatively, in case of parameter uncertainty, robust design optimization can be performed. Several methods and criteria exist for this purpose, and the method based on the HClnD-optimality criterion Φ_*HClnD*_ (eq. 3 is efficient [13], assuming an expected parameter distribution:

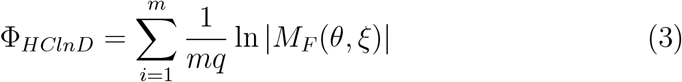

where there are *q* uncertain parameters and *m* = 2^*q*^. In this case *θ* are lower or upper percentile values of parameter distribution. In our study, we investigate robust designs according to this optimality criterion assuming 1 uncertain parameter (1UP or Robust 1UP) or 3 uncertain parameters (3UP or Robust 3UP).

In this study, optimizers are used to find the best allocation of sampling times, given *n*. The first category of algorithm performs optimization in a discrete design space. Discrete optimization requires defining the vector of all the possible sampling time allocations. We used the Fedorov-Wynn discrete algorithm [11, 28] implemented in PFIM version 4 [9] to find the optimal combination of sampling times for the illustrative studies of Mu/P1 and M13/P2, as well as for the phage ΦD9 decay experiment. The other category consists in algorithms working in a continuous design space, for which the sampling time window(s) and initial values for the design should be defined. We used the simplex continuous algorithm [20] implemented in PFIM version 4 to find the optimal sampling times according to viral parameters and the influence of design constraints (maximal duration and number of samples).

#### 2.1.3 Performance evaluation

The designs are compared in function of their optimality criteria Φ_*X*_ defined above. The X-efficiency *E*_*X*_ of a design *ξ*, with respect to a reference design *ξ*_*R*_ (for instance a non-optimized design), is computed as 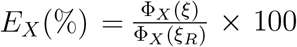. The expected standard errors of parameters are the main element of interest when evaluating a design. Relative standard errors (RSE, given in percentages, eq. 4) thresholds can be defined.

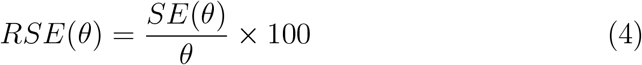

In this work, we consider that good RSE are below 30%, whereas poor and very poor RSE are greater or equal than 50% and 100%, respectively.

### 2.2 Application to ΦD9 multiphasic decay

#### 2.2.1 Viral decay experiment

For each replicate a separate phage stock was created by coculturing 2µL of frozen isogenic phage ΦD9 stock preserved at 80°C with 5 *×* 10^8^ ΔLamB ΔOmpC *E. coli* cells from an overnight culture into 10mL TrisLB (see Borin et al. [5] for media recipe and strain descriptions) at 37°C, 120rpm for 4 hours. The double receptor knockout host was used to select for the maintenance of OmpF use, which was previously found to be evolutionarily unstable. The relatively short incubation time was chosen to reduce the loss of fast decaying phage particles. Twelve phage lysates were filtered through 0.22 µm membranes and then a subsample of each stock was immediately plated to estimate initial viral titers. Plating was done on soft agar overlays with *E. coli* strain BW25113 and incubated overnight at 37°C [5]. Plaques were then enumerated. After initial phage samples were taken, 4mL of each lysate was then transferred to glass tubes that phages are not known to bind to [25] and incubated at 37°C and stationary to reduce evaporation. Additional phage samples were plated at 1, 2, 3, 5, 8, 24, 48, and 72 hours after the initial filtering.

#### 2.2.2 Data fitting and design optimization

Using the estimated parameters after data fitting from this experiment, robust design optimization were performed according to HClnD-criterion (3), supposing a possible triphasic decay, and assuming uncertain parameters for the two first phases and known parameters for the last phase and the initial viral density. The following design constraints were defined: a total of 9 sampling times including 0, 1, 2 and 3 days, and optimizing the 5 other sampling times among the vector (0.04, 0.08, 0.12, 0.17, 0.21, 0.25, 0.29, 0.33, 0.67, 0.71, 0.75, 0.79, 0.83, 0.88, 0.92, 0.96) d.

Population parameters were estimated by fitting the combined data from the replicates of an experiment, considering between-replicates differences as noise (the “naive pooled approach”) and following a maximum likelihood based method. Models were compared between them using the Akaike information criterion (AIC), which accounts for likelihood and the number of estimated parameters.

### 2.3 Software

R was used to perform design evaluation and optimization, through the PFIM version 4.0. program [9] and figures. Data fitting was performed with Monolix version 2021R2 [17]. Codes for design optimization and data fitting are available in Supplementary files.

### 2.4 Data availability

The data sets used and/or analyzed during the current study are available from the corresponding author on reasonable request. The codes to perform design evaluation and optimization, as well as data fitting are accessible at: https://github.com/jeremyseurat/DesignMultiphasicViralDecay.

## 3. RESULTS

### 3.1 Impact of multiphasic rates of population decay - concept

The decay of virus populations is impacted by the number of ‘phases’, corresponding decay constants, and the proportions of each subpopulation in the population as a whole. To explore these effects, we illustrate the impact of biphasic decay on total virus populations when modulating the associated decay rate and proportion of subpopulations. In case of a monophasic decay, the viral load (*V*) decays over time (*t*) as follows (eq. 5):

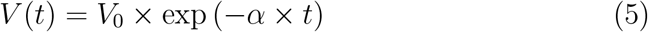

with *V*_0_ the initial viral density which could be estimated or known from the study settings and *α* the decay rate. In this case, as long as viral density is quantifiable, a limited number of measurements (e.g., two or three) over time are sufficient to estimate *α* with adequate precision. In the case of biphasic decay, the viral load decays according to the following:

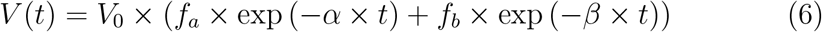

with the parameters *f*_*a*_ the initial proportion of the phage of decay rate *α* and *f*_*b*_ for the phage of decay rate *β, f*_*a*_ = 1 − *f*_*b*_. The impact of the different parameters (decay rates and initial proportions) on the viral kinetics is shown in Figure 1. The two decay phases can be distinguished in the case of two very different rates, for example *α* = 0.5 /d and *β* = 0.05 /d with similar proportions of the two subpopulations. The two slopes could also be readily observable if the fast decaying subpopulation represents a much larger fraction than the slow decaying subpopulation (e.g., *α* = 0.1 /d, *β* = 0.5 /d, *f*_*a*_ = 0.05 and *f*_*b*_ = 0.95). However, distinguishing the two phases is more complex when decay rate values are close to each other or when the slowestdeclining viral subpopulation represents a large proportion of the total.

**Figure 1:**
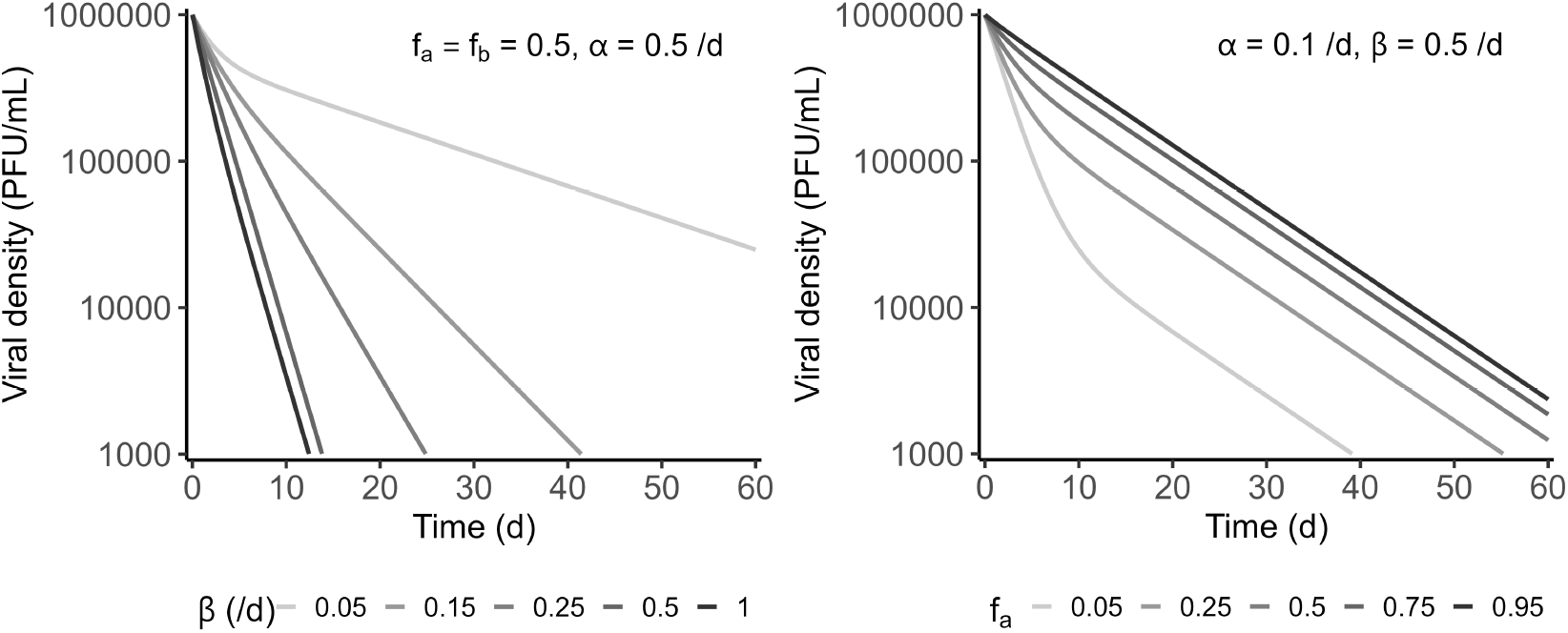
Influence of the viral parameters on a biphasic decay. Biphasic decays (eq. 6) are represented on the two panels: with different decay rates for one population (*β*) ranging from 0.05 to 1 /d (with same initial proportions for both populations i.e., *f*_*a*_=*f*_*b*_=0.5 and decay rate of the other population (*α*) of 0.35 /d) on the left and different initial proportions *f*_*a*_ ranging from 0.05 to 0.95 (with *f*_*b*_=1-*f*_*a*_, *α*=0.1 /d and *β*=0.3/d) on the right.

### 3.2 Decay rates, inference and design optimization: illustrative example

To illustrate how the viral decay rates, and the design (measurement times) choice can influence the precision of parameter estimation, we choose two different synthetic examples based on measurements of two phage with known decay rates when isolated at 37 °C [7].

In the first example, we consider a study composed of two phages with very different decay rates such as Mu (with a decay rate of 0.29 /d) and P1 (with a decay rate of 0.077 /d). We assume similar proportions of the two phages (*f*_*a*_=*f*_*b*_=0.5), and an error parameter (notably reflecting the noise) *σ* of 0.2 (i.e., 20% standard deviation of the discrepancy between data and the true viral density). Using a non-optimized design with 7 equispaced sampling times (1 sample every 10 days), the precision of parameters relative to the fastest phage, Mu are poor with a relative standard error (RSE) of 101% for the estimation of the decay rate of Mu, which is due to the lack of information in the early phase of the decay kinetics.

For the same viruses, if we fix the initial sampling time (t=0 d) and optimize the 6 other sampling times (among (2, 4, 6, 8, 10, 12, 14, 16, 18, 20, 22, 24, 27, 30, 35, 40, 50, 60) d), the optimal design is composed of the following sampling times: (0, 4, 6, 20, 22, 50, 60) d. This design allows a better overall precision of parameters (gain in D-efficiency of 24%, see Methods) and the RSE of the decay rate of Mu is expected to be 63% (vs. 101% without design optimization, Table 1). However, if the two phages have a similar decay rate, for instance M13 (0.074 per day) and P2 (0.041 per day), the expected precision of the different parameters is very poor (RSE*>*100%) even using an optimized design.

**Table 1:**
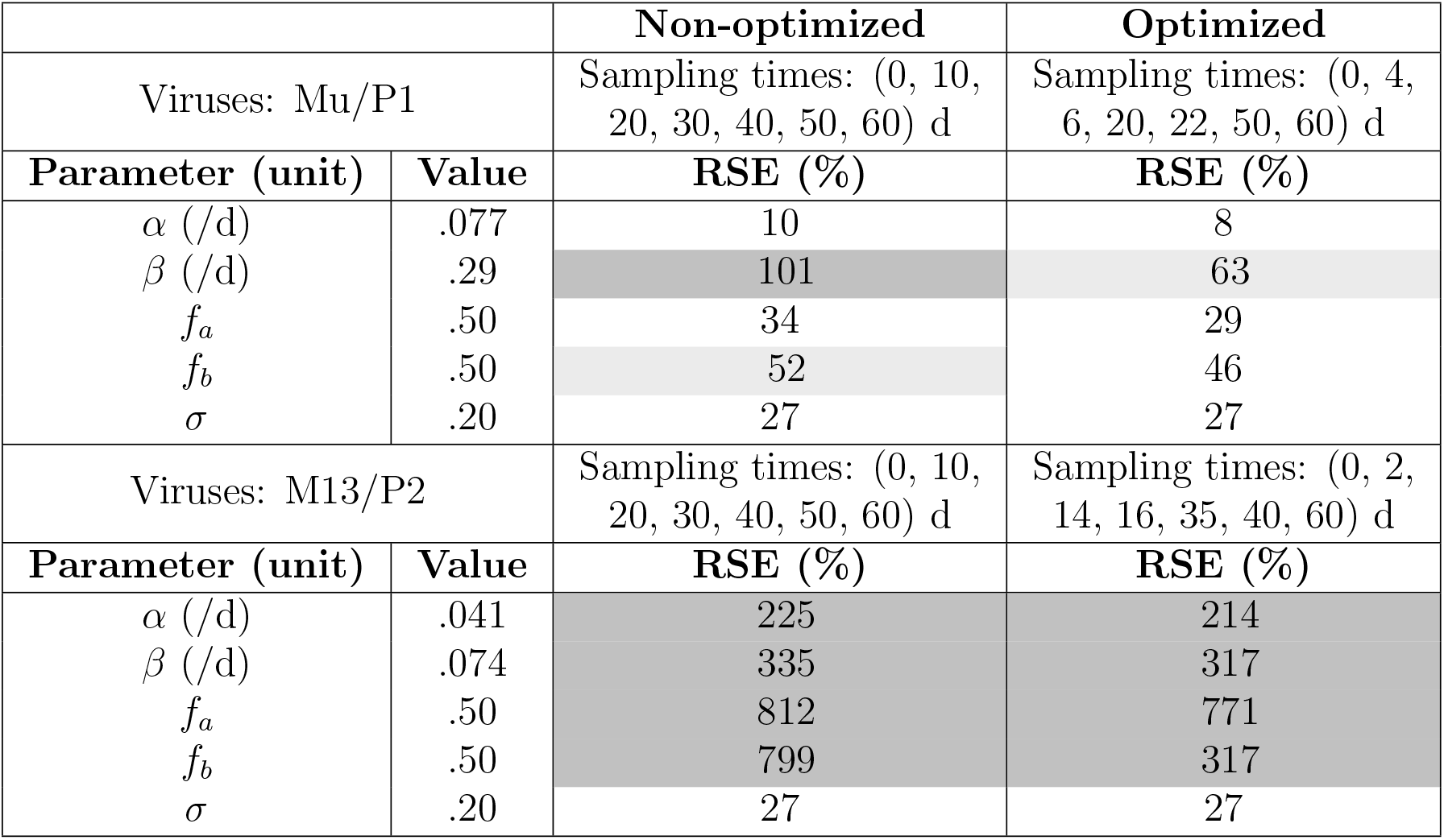
Viral decay rates effect on expected precision of estimates from a nonoptimized or optimized design. The phage subpopulations Mu and P1 are represented in the top part and M13 and P2 in the bottom part. The vector of sampling times is given in days (d), *α* is the decay rate of the slowest virus (Mu and M13, respectively), *β* is the decay rate of the fastest virus (P1 and P2, respectively), *f*_*a*_ and *f*_*b*_ are the initial proportions (corresponding to *α* and *β*, respectively), *σ* is the proportional error parameter, RSE is the relative standard error: a RSE greater than 50% or 100% is indicated by a light grey or a dark grey cell, respectively.

|  |  | Non-optimized | Optimized |
| --- | --- | --- | --- |
| Viruses: Mu/P1 |  | Sampling times: (0, 10, 20, 30, 40, 50, 60) d | Sampling times: (0, 4, 6, 20, 22, 50, 60) d |
| Parameter (unit) | Value | RSE (%) | RSE (%) |
| $\alpha$ (/d) | .077 | 10 | 8 |
| $\beta$ (/d) | .29 | 101 | 63 |
| $f_a$ | .50 | 34 | 29 |
| $f_b$ | .50 | 52 | 46 |
| $\sigma$ | .20 | 27 | 27 |
| Viruses: M13/P2 |  | Sampling times: (0, 10, 20, 30, 40, 50, 60) d | Sampling times: (0, 2, 14, 16, 35, 40, 60) d |
| Parameter (unit) | Value | RSE (%) | RSE (%) |
| $\alpha$ (/d) | .041 | 225 | 214 |
| $\beta$ (/d) | .074 | 335 | 317 |
| $f_a$ | .50 | 812 | 771 |
| $f_b$ | .50 | 799 | 317 |
| $\sigma$ | .20 | 27 | 27 |

### 3.3 Optimal time measurements according to viral parameters

Next, we studied more extensively the influence of the decay rate of the two viral subpopulations on the RSE of resulting fits (different combinations of decay rates between 0.02 and 0.5 /d, Figure 2). Assuming equal initial proportions of the two viruses, a non-optimized design (1 sampling time every 10 days during 60 days) leads to poor RSE and potentially the nonidentifiability of parameters if *β* (the fastest decay rate) is higher than 0.35 /d and *α* (the slowest) is lower than 0.15 /d or higher than 0.3 /d. An optimal design composed of the same number of sampling times found using the simplex algorithm (i.e., finding the 7 optimal times in the continuous design space [0,60] d) can provide acceptable RSE in most of those days. However, as in the example shown in Table 1, two very close decay rates for instance which differ from 0.06 /d are difficult to estimate properly, even with design optimization.

**Figure 2:**
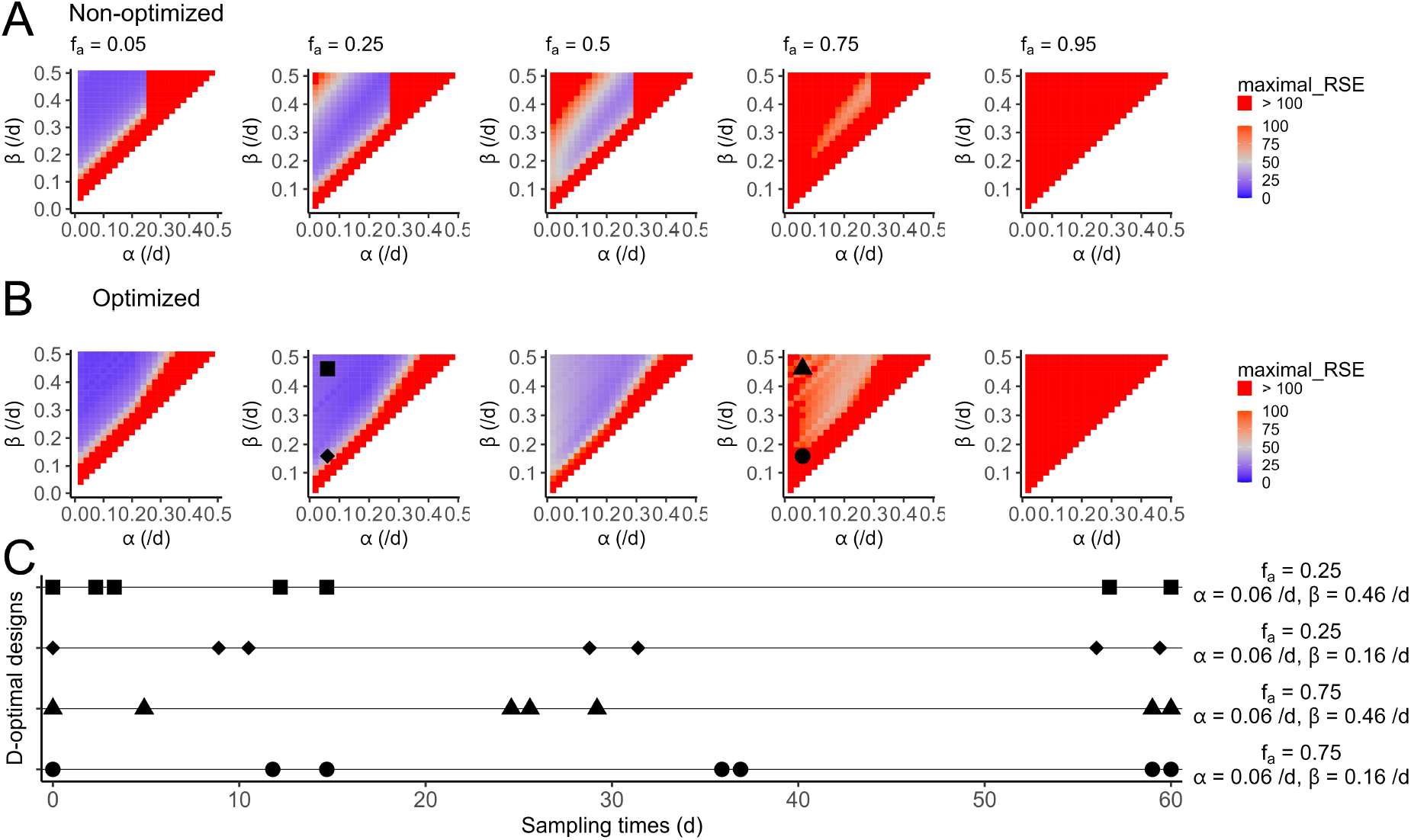
Influence of viral parameters on maximal imprecision (RSE) of estimates using an (A) empirical non-optimized design or an (B) optimized design and on (C) optimal sampling times. The non-optimized design consists in 7 measuring times from 0 to 60 days every 10 days. The design optimization is performed by the simplex algorithm (7 times in the [0-60] days window), where *α* is the decay rate of the slowest virus, *β* is the decay rate of the fastest virus, *f*_*a*_ and *f*_*b*_=1-*f*_*a*_ are the initial proportions (corresponding to *α* and *β*, respectively). RSE refers to relative standard error and the color represents the higher expected RSE among the estimated parameters: *α, β, f*_*a*_, *f*_*b*_ and *σ* (the error model parameter). Optimal designs are represented for different parameters combinations by specific symbols (*α* = 0.06 /d, square: *β*=0.46/d and *f*_*a*_=0.25, rotated square: *β*=0.16/d and *f*_*a*_=0.25, triangle: *β*=0.46/d and *f*_*a*_=0.75, circle: *β*=0.16/d and *f*_*a*_=0.75).

From the influence of the initial proportion of the two viral subpopulations *f*_*a*_ and *f*_*b*_ (Figure 1), we understand that the identification of the two phases of the decay is easier with a smaller proportion of viruses that have the slower decay rate. This is confirmed by Figure 2 where the maximal RSE is never below 100% if *f*_*a*_=0.95 regardless of *α* and *β* values between 0.02 and 0.5 /d, even if the 7 measuring times are optimized in the [0,60] days window.

However, design optimization can improve the precision of estimation when the initial proportions are more balanced (*f*_*a*_ between 0.25 and 0.75), especially when one of the decay rates is much faster than the other one. Interestingly, the optimization algorithm tends to identify optimal measurement times earlier when one of the two decay rates is high than when both decay phases are slow (e.g., *α* = 0.06 /d, *β* = 0.46 /d vs. *α* = 0.06 /d, *β* = 0.16 /d, Figure 2). In the case of very low initial proportion of the slowest-decaying virus (*f*_*a*_=0.05), a non-optimized design is generally good enough to provide sufficient RSE in most of the cases.

### 3.4 Influence of design constraints

Another possibility to reach sufficient information from the experimental design is to increase the number of sampling times during the study. In this section, we evaluate how many measurements would be needed to achieve precise parameter estimates (RSE *<* 30%), depending on the decay rates of the two populations which are assumed to be in the same initial proportions (i.e., *f*_*a*_ = *f*_*b*_ = 0.5). In doing so, we investigate if considering longer studies (120 days instead of 60 days) can help to quantify precisely the two slopes during biphasic decay in this particular case study.

Increasing the number of measurements increases the quantity of information, and as a consequence improves the precision of parameters. For instance, in the case of *α* = 0.1 /d and *β* = 0.4 /d, 7 optimized samplings during the 60 days following the beginning of the study are not sufficient to obtain RSE *<* 30% in all parameters. In this specific case, 10 sampling times are needed to reach this objective (Figure 3). This option however increases the cost of the study. Furthermore, optimizing sampling times may already provide sufficient information. In our example, there are cases where 5 measuring times would be sufficient to obtain precise estimates, for example if *α* = 0.24 /d and *β* = 0.44 /d.

**Figure 3:**
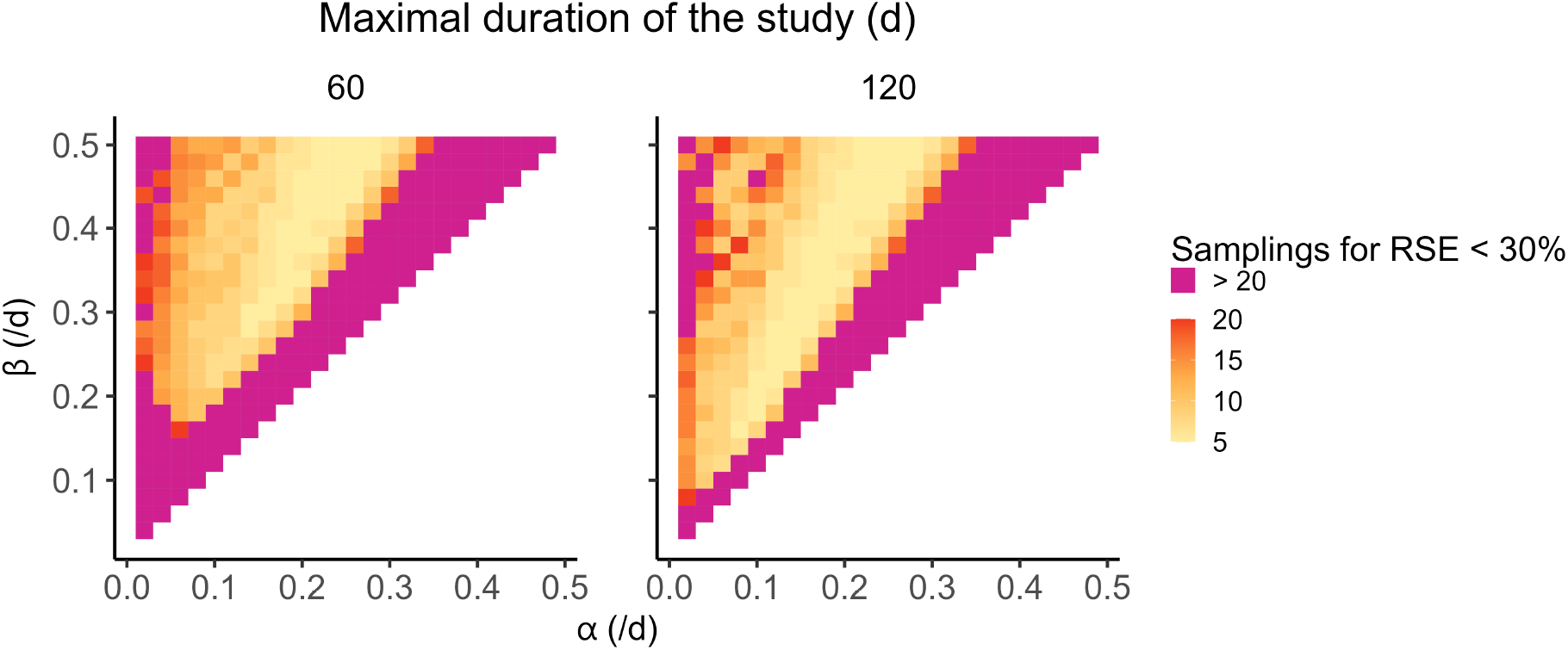
Influence of maximal duration time on needed sampling times to reach adequate precision of estimation. A flexible number of measuring times is considered between 0 and 60 days on the left and between 0 and 120 days on the right, respectively. Sampling times of the design are optimized using the simplex algorithm in the corresponding sampling window ([0,60] and [0,120]). *α* is the decay rate of the slowest virus, *β* is the decay rate of the fastest virus, the initial proportion of each viral population is 0.5. RSE: Relative Standard Error.

The duration of the study is another important design consideration which depends on the two decay rates. For example when *α* is 0.1 /d and *β* is 0.2 /d, allowing a maximal duration of 120 days instead of 60 days can reduce the number of measurements by a factor 2 (5 vs. 10, respectively) to reach the RSE objective. Importantly, extending the duration of the study is needed to investigate the decay of two viruses with slow elimination: for instance if *α* is 0.08 /d and *β* = 0.12/d, choosing a maximal duration of 120 days enables precise parameter estimation, which is not the case with a 60day study design, regardless of the number or the allocation of the sampling times. In constrast, the design can be shortened if we know that one of the two populations have a fast decay rate. Indeed, similar number of sampling times are needed between 60 and 120 days when *β* is between 0.4 and 0.5 /d (Figure 3).

### 3.5 Finding optimal design accounting for parameter uncertainty

In many cases, we do not have initial estimates of the parameters associated with the loss of infectious virus particles over time. A possibility is to assume the uncertainty on one or several parameters, and define a prior distribution. Robust designs account for the parameter uncertainty during the optimization procedure. Defining priors based on the distribution of viral decays in [7], we investigated two robust design scenarios accounting for uncertainty. In the first scenario, one viral decay rate was uncertain (between 0.04 and 0.35 /d, the 5th and 95th percentiles of the decay rates in [7], respectively), i.e., 1 Uncertain Parameter (1UP), the other one was set at 0.09 /d (the median decay rate in [7]) and initial proportions were 50% for the two populations (*f*_*a*_=*f*_*b*_=0.5). In the second scenario, we considered uncertainty on all the parameters, i.e., 3 Uncertain Parameters (3UP): the two viral decay rates (between 0.04 and 0.35 /d and between 0.06 and 0.19 /d, respectively) and the initial proportions (between 0.05 and 0.95) for each population.

Robust design optimization is performed following the FIM based HClnDcriterion (see Methods), following a combinatorial procedure over the 27132 possible designs of 7 sampling times. The initial time (t = 0 d) is included in each possible design and the 6 other samplings are optimized among the following vector of possible times: (1, 2, 3, 4, 5, 6, 8, 10, 13, 16, 20, 24, 28, 32, 36, 40, 45, 50, 60) d.

The allocation of sampling times for the robust designs was (0, 4, 5, 16, 20, 40, 60) days in the case of 1 parameter uncertainty and (0, 4, 10, 20, 32, 50, 60) days in the 3 parameter uncertainty scenario. We compare these methods and scenarios with two other design procedures: a non-optimized (equispaced) design and the local optimal design (i.e., optimal design if we knew both decay rates and the subpopulation fractions). Accounting for the different parameter possibilities (decay rates between 0.02 and 0.5 /d, consistent with the data in [7], and initial proportions between 0.05 and 0.95), the coverage is defined as the proportion of cases where a certain precision of estimate is reached, i.e., a RSE *<* 30% or 50% in function of the design strategy (Figure 4). The detailed precision of estimates with robust designs for each possible parameter combination are provided in Figure S1.

**Figure 4:**
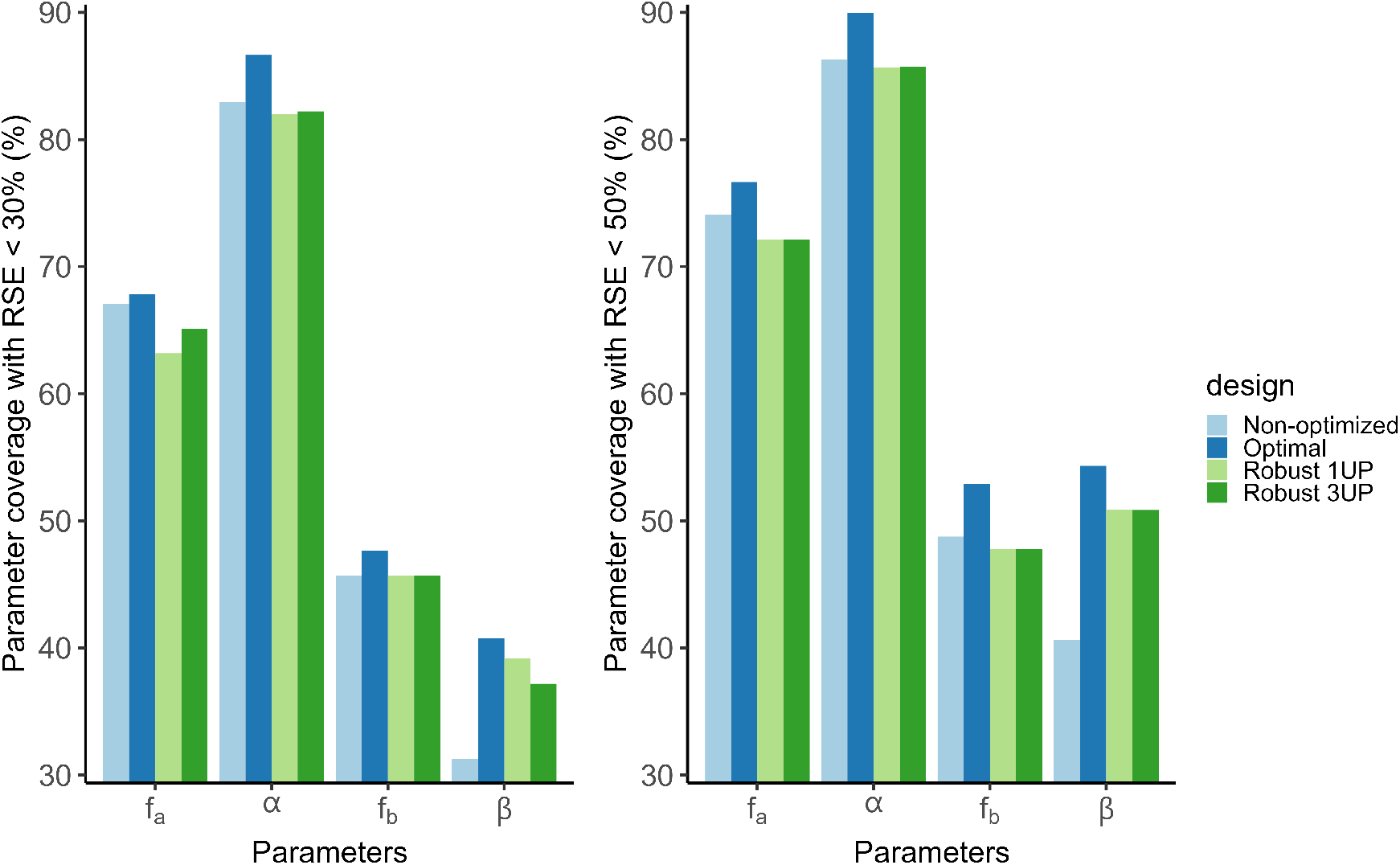
Design comparison for expected coverage of parameter precision. Coverage is the percentage of parameter settings (*α* from 0.02 to 0.48 /d (increment of 0.02), *β*(*> α*) from 0.04 to 0.5 /d, *f*_*a*_ and *f*_*b*_ among (0.05, 0.25, 0.5, 0.75 or 0.95, with *f*_*a*_+*f*_*b*_=1) that the different design strategies allow to reach an expected RSE (Relative Standard Error) of 30% (left) or 50% (right) for the different parameters.

Parameters relative to the virus with faster elimination (*f*_*b*_ and *β*) were the most difficult to estimate precisely. For instance, reaching a good RSE of 30% or acceptable RSE of 50% on the estimation of *β* was covered in respectively 31% or 41% of cases if the design was not optimized. Due to early time points (5 in the first 20 days for Robust 1UP and 4 for Robust 3UP vs. 1 point every 10 days in the non-optimized design), the robust designs allowed to reach coverages of 39% (1UP) and 37% (3UP) or 51% (1UP and 3UP), respectively. Note that the performances of robust designs in case of parameter uncertainties are close to the coverage values of optimal designs (41% with RSE below 30% or 54% with RSE below 50%) if we were to know the parameters at the design step. In contrast, coverages were better and comparable regardless of the design for the estimation precision of parameters relative to the slower virus (*f*_*a*_ and *α*). The key point is that optimal design was able to significantly improve the fraction of contexts in which the fast decay rate *β* was precisely estimated compared to non-optimized designs and even in the face of parameter uncertainty (see right-most columns, in Figure 4).

### 3.6 Application: design optimization of phage ΦD9 decay

A pilot experiment of phage ΦD9 decay was performed using a nonoptimized empirical design of 7 days with one measurement each day (every 24 h) except at day 6. The data showed evidence of a multiphasic decay (Figure 5A). However, data fitting with the biexponential model (eq. 6, in that case, *V*_0_ and *f*_*a*_ were estimated given and *f*_*b*_=1-*f*_*a*_) showed that a precise estimation of the first (fastest) phase of this decay (*β*) was not possible (Table 2). The lack of precision arose because the rapid decay of the fastest subpopulation appeared to occur during the first day without sufficient sampling.

**Table 2:** Parameter estimates from the different designs of ΦD9 decay. The vector of sampling times is given in days (d), *α* is the decay rate of the slowest virus subpopulation, *β* is the decay rate of the fastest subpopulation, *V*_0_ is the initial total virus density, *f*_*a*_ and *f*_*b*_ are the initial proportions (corresponding to *α* and *β*, respectively), *σ* is the proportional error parameter, RSE is the relative standard error: a RSE greater than 100% is indicated by dark grey cell, and hyphen is indicated if the parameter was fixed (i.e., not estimated).

|  | **Non-optimized**<br>Sampling times: (0, 1, 2, 3,<br>4, 5, 7) d |  | **Optimized**<br>(0, .04, .08, .13,<br>.21, .33, 1, 2, 3) d |  |
| --- | --- | --- | --- | --- |
| **Parameter (unit)** | **Estimate** | **RSE(%)** | **Estimate** | **RSE(%)** |
| $\alpha$ (/d) | .18 | 25 | .19 | 36 |
| $\beta$ (/d) | 39 | NaN | 3.2 | 6 |
| $V_0$ | $3.8 \cdot 10^4$ | 20 | $6.1 \cdot 10^4$ | 3 |
| $f_a$ | .17 | 28 | .06 | 17 |
| $f_b (=1-f_a)$ | .83 | - | .94 | - |
| $\sigma$ | .70 | 8 | .17 | 7 |

**Figure 5:**
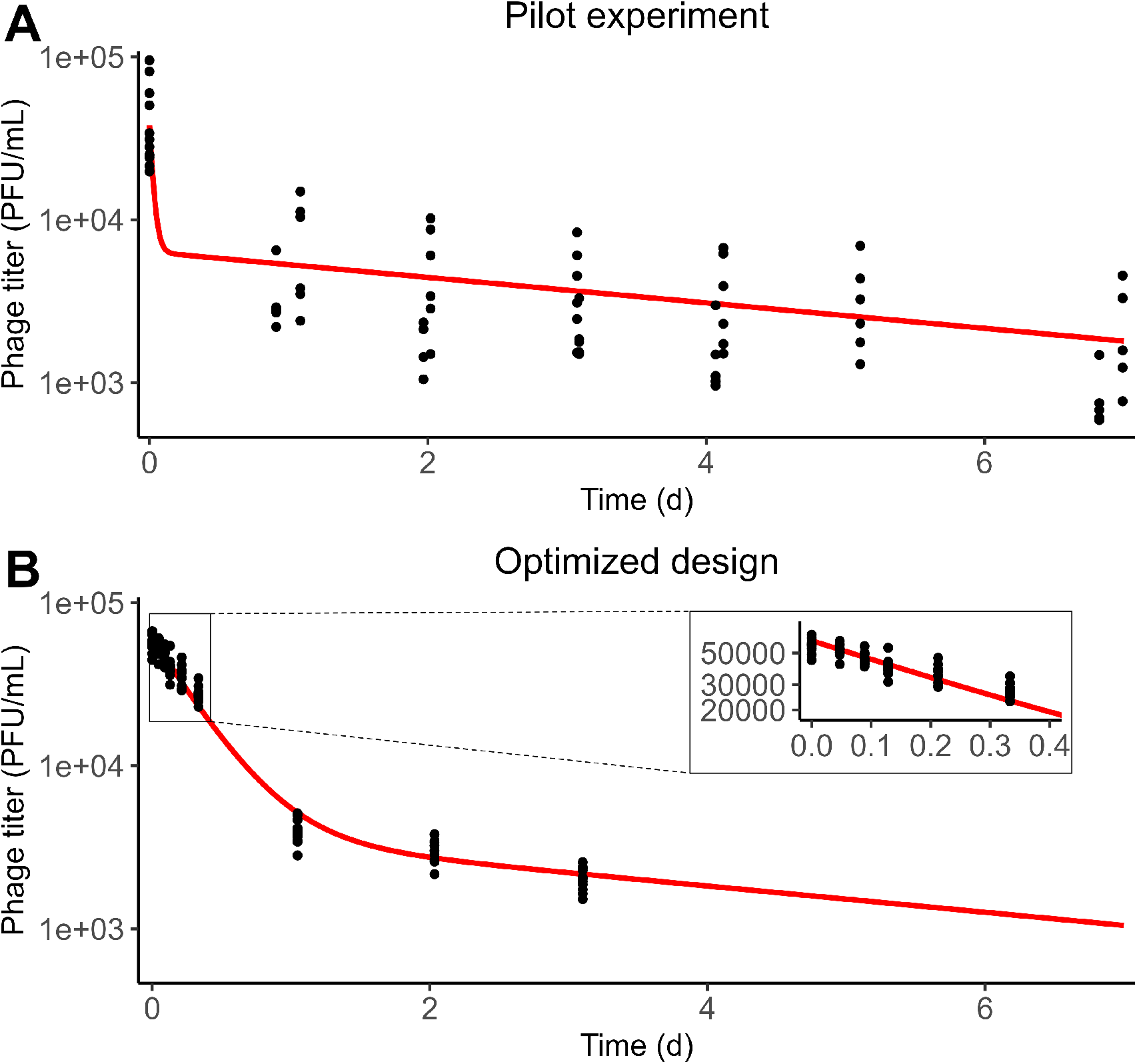
Phage ΦD9 decay data fitting from (A) the non-optimized pilot experiment and (B) the optimized design. Red line is the viral density decay prediction from the biexponential model (eq. 6) and the parameters given in Table 2. The pilot experiment was composed of two batches with theoretical sampling times at 0, 1, 2, 4, 5, and 7 days for one batch and 0, 1, 2, 4, and 7 days for the other batch. Six replicates were made for each measurement of the pilot experiment. The optimized design experiment was composed of one batch with theoretical sampling times at 0, 0.04, 0.08, 0.13, 0.21, 0.33, 1, 2, and 3 days (i.e., 0, 1, 2, 3, 5, 8, 24, 48, and 72 hours). Twelve replicates were made for each measurement of the optimized experiment.

Therefore, our design aim was to propose revised sampling times for a follow-up experiment allowing a precise estimation of the fast decay rate *β* and other parameters both the slow decay rate *α* and the fraction of subpopulations, *f*_*a*_ = 1 − *f*_*b*_. An additional objective was to propose a design that could allow to identify whether there was triphasic decay, comprised of two highly unstable subpopulations that both decayed during the first day. Given the knowledge from the first experiment on the slower subpopulation and its initial density, we defined the following design constraints: a total of 9 sampling times including 0, 1, 2 and 3 days, and optimizing the 5 other sampling times located between 0 and 1 day.

The triphasic decay model was written as follows (eq. 7):

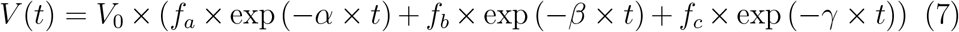

with *f*_*a*_, *f*_*b*_, and *f*_*c*_ the initial proportions of the viral subpopulations of decay rates of *α, β* and *γ*, respectively. Since biphasic and triphasic decay are nested models (contrast eq. 6 with eq. 7), robust design optimization was performed for the most complex model, i.e., the triphasic decay model, assuming that a design can estimate three phases of viral decay is also adequate to estimate two phases if only two are observable. From the inference of the pilot experiment, we assumed *α, f*_*a*_ and *V*_0_ as well-constrained prior parameters (see values in Table 2). Using the HClnD-criterion based optimization (see Methods), we accounted for uncertainty on the parameters relative to the two viral subpopulations with fastest rate. The robust design was composed of the following sampling times: 0, 0.04, 0.08, 0.13, 0.21, 0.33, 1, 2 and 3 days.

Data collected from this second experiment, with optimized sampling times, enables precise estimates of the parameters of the biphasic decay model (eq. 6), including the faster decay phase, despite a higher initial density (Figure 5B, Table 2). Indeed, the first decay rate *β* was mis-estimated during the pilot experiment as a very high value (39 /d with RSE = NaN). In contrast, using the optimized design in the second experiment, we identified and estimated *β* as 3.2 /d, with a high precision (RSE = 6%). The estimation of the slower decay rate *α* was re-estimated to 0.19 /d with a moderate precision (RSE = 36%), which is consistent with the value obtained from the first experiment of 0.18 /d. We can also observe a lower noise than expected (the error parameter *σ* was equal to 0.17 compared to 0.70 from the fitting of the first experiment).

A triphasic model (eq. 7) with a faster initial decay did not improve the fit (Figure S2): AIC was lower for the biphasic model: 2045.7 vs. 2049.7 for the triphasic model, i.e., the biphasic model is more parsimonious. Even though our experiment included 7 samplings on the first day, we were unable to detect more than two subpopulations (Table S1). It is possible that multiple subpopulations exist with similar decay rates, however alternative approaches would be required to identify them.

## 4. DISCUSSION

Here we revisited classic assumptions of exponential decay of virus particles to evaluate, optimize, and implement an alternative approach to measuring virus densities over time. In doing so, we show when and how an information-theoretic method can be used to leverage prior information and construct practical experimental designs that can accurately infer the proportion of viral subpopulations and their corresponding decay rates. We then successfully implemented an information-theoretic design to infer both a slow and fast decay rate in phage ΦD9, outperforming conventional equispaced design schemes.

The implementation of an information theoretic design scheme provides a route to distinguish biphasic from monophasic decay. However, we recognize that there are limits to such inference given uncertainty in the underlying decay model. Alternative methods exist to perform design optimization in the case of structural model uncertainty [18, 26]. In our real application, data from the two experiments were fitted separately, the first one being only used to design the second experiment. One potential alternative would have been to pool the data from multiple experiments to improve parameter precision of estimation, as in model-based adaptive designs [23, 10]. Nonetheless, we leveraged the first experiment to establish priors accurately estimated parameters and not for those with significant uncertainty - modulated in part by the impact of noise on inference. We caution that noise has a linear impact on parameter imprecision such that measurement error propagates linearly to impact RSE - a significant issue in estimating rapid decay.

In this study, subpopulations of phage are initially defined using models with a single decay rate corresponding to each subpopulation. In reality, complex processes of phage-bacteria co-evolution can give rise to many subpopulations [22, 5]. Moreover, it is also possible that other epigenetic and/or biophysical mechanisms could lead to multiphasic decay. For example, aggregation of virus particles may lead to faster viral decay at high virion concentrations, with models including quadratic decay relevant to representing this phenomenon [3]. In addition, it is well known that experimental context, including temperature, can influence virion stability and, in turn, the decay of infectious virus particles [15, 7]. In the present context, we cannot distinguish between the underlying mechanisms. We are optimistic that using information theoretic methods as part of experimental design can be used to identify the generality and relevance of multiphasic decay for phage and other viruses, as has been shown for HIV [27], and in bacterial systems [6].

Moving forward, alternative optimal experiment design methods could be used to identify the relevance of multiphasic decay in viral populations. These experimental designs could include ElnD-criterion based robust designs which account for the expected distribution of uncertain parameters but are more time-consuming to compute relative to than HClnD-designs [13]. In such cases, the design choice can also be influenced by priorities, e.g., choosing to focus on estimating specific parameters of a biphasic decay, such as the fastest decay rate. For this purpose, the *DD*_*S*_-criterion can be used at it allows an optimal balance between the parameter(s) of interest and other parameters of the models [2]. A cost function accounting for the number of samples can also be included during design optimization. Finally, alternative optimization algorithms may also be appropriate, notably when trying to infer parameters associated with decay in the context of more complex models [29, 16]. If optimal design theory is not applicable, one option could be to utilize equispaced sampling times on a log-scale rather than linear-scale after initial prototypes are used to establish a relevant time-scale for decay.

To conclude, this study propose methods to design viral decay experiments and shows the conceptual and practical value of design evaluation and optimization. By leveraging prior information and a pilot experiment, we were able to design an improved, follow-up experiment that robustly inferred the presence of a large majority of fast-decaying viruses within the phage ΦD9 population. Beyond the practical value of improving specific parameter estimates, the use of robust design may also open the door to identifying the genetic and/or epigenetic basis for variation in viral life history traits.

## 5. FUNDING

This work was supported by the Howard Hughes Medical Institute Emerging Pathogens Initiative grant 7012574 (J.R.M.), National Institutes of Health grants T32GM007240 (J.M.B.) and 1R01AI46592-01 (J.S.W.), and the Chaire Blaise Pascal program of the Île-de-France region (J.S.W.).

## 6. AUTHOR CONTRIBUTIONS

JS: Conceptualization, Methodology, Software, Formal analysis, Data Curation, Writing - Original Draft, Visualization. KRG: Investigation, Resources, Writing - Original Draft. JRM: Validation, Resources, Writing - Review & Editing, Supervision, Funding acquisition. JMB: Validation, In- vestigation, Writing - Review & Editing Resources, Supervision, Funding acquisition. JSW: Conceptualization, Validation, Writing - Review & Editing, Supervision, Funding acquisition.

## 7. COMPETING INTERESTS

The authors have no competing interest to declare.

## Notes

### Competing Interest Statement

The authors have declared no competing interest.

### Summary of Updates

Clarifications are provided to ease the understanding the manuscript. One word of the title has been updated as well.

