## Supplementary material for "Design, optimization, and inference of biphasic decay of infectious virus particles"

---

---

| | Biphasic model | | Triphasic model | |
|---|---|---|---|---|
| **Parameter (unit)** | **Estimate** | **RSE(%)** | **Estimate** | **RSE(%)** |
| $\alpha$ (/d) | .19 | 36 | .19 | 52 |
| $\beta$ (/d) | 3.2 | 6 | 3.2 | >1000 |
| $\gamma$ (/d) | - | - | 3.2 | >1000 |
| $V_0$ | 6.1 $10^4$ | 3 | 6.1 $10^4$ | 3 |
| $f_a$ | .06 | 17 | .06 | 28 |
| $f_b$ | .94 | - | .07 | NaN |
| $f_c$ | - | - | .87 | - |
| $\sigma$ | .17 | 7 | .17 | 7 |

**Table S1**: **Parameter estimates from the different models and ΦD9 decay.** The decay rate of the slowest virus subpopulation is denoted $\alpha$., whereas $\beta$ and $\gamma$ are the decay rate of the fastest subpopulations. $V_0$ is the initial total virus density, $f_a$, $f_b$ and $f_c$ are the initial proportions (corresponding to $\alpha$, $\beta$ and $\gamma$, respectively), $\sigma$ is the proportional error parameter, RSE is the relative standard error: RSE greater than 50% or 100% are indicated by light grey or dark grey cells, respectively, and hyphen is indicated for $\gamma$ and $f_c$ in case of biphasic model as non-existing parameters and in RSE column if the parameter was fixed (i.e. not estimated).

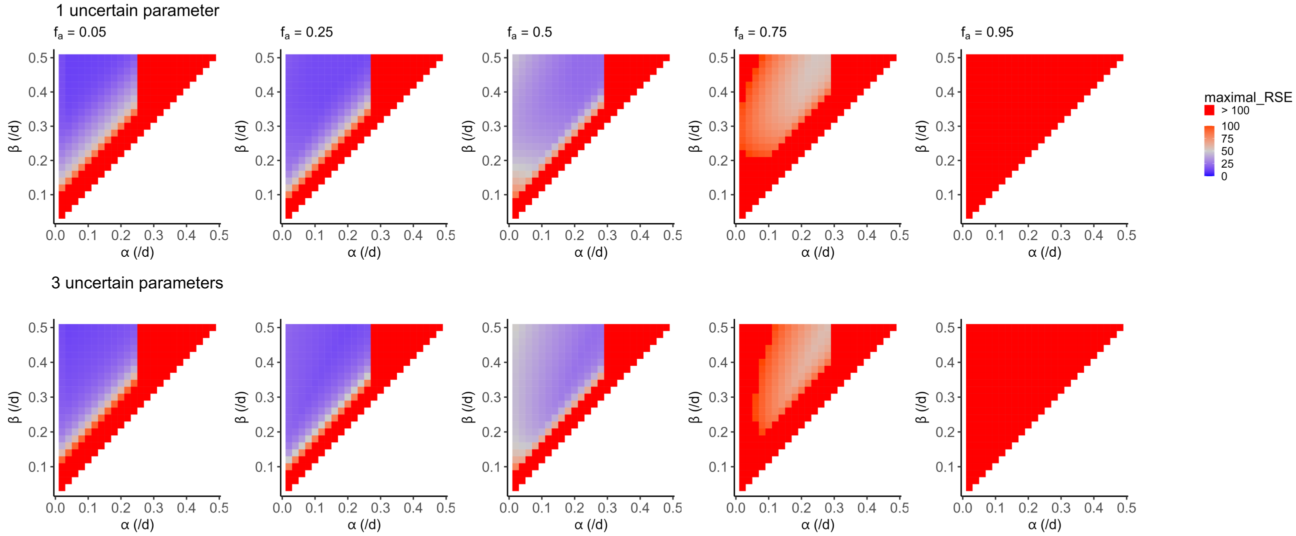


**Figure S1**: **Robust design optimization accounting for parameter uncertainties.** Designs are optimized according to HClnD criterion. In the first row, we account for uncertainty on one of the two viral decay rates $\alpha$ and assume known $\beta$=0.09 /d and initial proportions of each virus ($f_a$=$f_b$=0.5). In the second row,we account for uncertainty on the two viral decay rates and on the initial proportions of each virus. The design are obtained by combinatorial optimization among 27132 designs of 7 sampling times in the [0-60] days window. RSE refers to relative standard error and the color represents the higher expected RSE among the estimated parameters: $\alpha$, $\beta$, $f_a$, $f_b$ and $\sigma$ (the error model parameter).

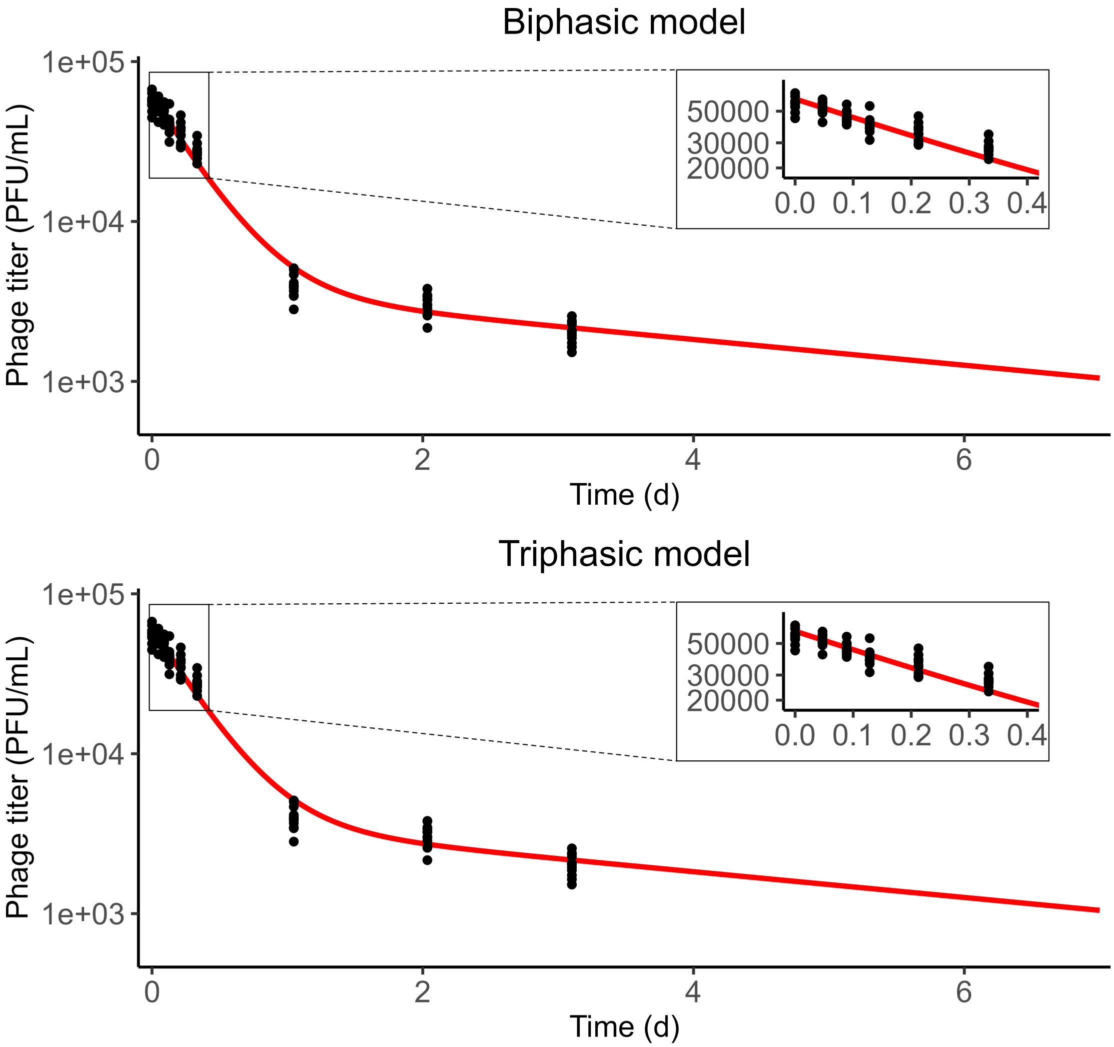


**Figure S2**: **Phage ΦD9 decay data fitting using a biphasic decay (top) or a triphasic decay (bottom).** The line is the viral density decay prediction from the biexponential model (eq. 6) or from the triexponential model (eq. 7) and the parameters given in the Table **S1**)
